# Distinct microbial communities in the murine gut are revealed by taxonomy-independent phylogenetic random forests

**DOI:** 10.1101/790923

**Authors:** Gurdeep Singh, Andrew Brass, Sheena M. Cruickshank, Christopher G. Knight

**Author notes:** **Corresponding author:** Sheena M. Cruickshank, A.V. Hill Building, The University of Manchester, Oxford Road, Manchester, M13 9PT.

## Abstract

Gut microbiome analysis using 16S rRNA frequently focuses on summary statistics (e.g. diversity) or single taxonomic scales (e.g. Operational Taxonomic units, OTUs). This approach risks misinterpreting the phylogenetic or abundance scales of community differences (e.g. over-emphasising the role of single strains). We therefore constructed a 16S phylogenetic tree from mouse stool and colonic mucus communities. Random forest models, of all 428,234 clades, tested community differences among niches (stool versus mucus), host ages (6 versus 18 weeks), genotypes (wildtype versus colitis prone-*mdr1a^-/-^*) and social groups (co-housed siblings). Models discriminated all criteria *except* host genotype, where no community differences were found. Host social groups differed in abundant, low-level, taxa whereas intermediate phylogenetic and abundance scales distinguished ages and niches. Thus, treating evolutionary clades of microbes equivalently without reference to OTUs or taxonomy, clearly identifies whether and how gut microbial communities are distinct and provides a novel way to define functionally important bacteria.

## Introduction

Microbiome studies can be inaccurate or biased for both experimental and analytical reasons. Analytically, there is often a focus on how changes in individual taxonomic levels impact upon the host. For instance, large-scale shifts among phyla (notably the Firmicutes to Bacteroidetes ratio (1)) and particular species differences (*Lactobacillus reuteri* enrichment (2)) have each been associated with obesity. However bacteria cooperate within communities and thus the power of such individual statistics, at whatever level, to distinguish the complexities of communities is extremely limited. For example, it may lead to bias and over-interpretation of the relative importance of single species changes. Even if a more complete phylogenetic tree is considered, it is often relegated to calculating individual diversity statistics by UniFrac (3). The value of such metrics in the face of technical variation among amplicon sequencing protocols and their implementation in different laboratories is also questionable (4). To date, it is rare to examine taxa at different levels in a single analytical framework, enabling the relative importance of community differences at different phylogenetic scales to be assessed.

Even where studies consider multiple phylogenetic levels together (5), analyses are highly constrained by available taxonomies – even the addition of operational taxonomic units (OTUs) to the traditional Linnaean hierarchy from domains to species does not come close to capturing the diverse intricacy of microbial phylogeny (6). Such taxonomies are inadequate, not only because all taxonomies are incomplete and different taxonomies disagree (7, 8), but because only a small amount of the evolutionary history of any group of microbes can be captured in a taxonomy with a limited number of levels. A risk of bias also applies to the widely-used, but necessarily arbitrary, 97% or 99% similarity thresholds for grouping sequences into OTUs (9). While it may represent a biologically-based compromise, no such threshold can work equally well across the bacterial tree (10).

Bias may also stem from experimental design issues, such as the sampling site of the microbiota and the environment of the host. For instance, gut microbiota research has tended to focus on stool samples. Stool samples have shown changes in the microbiota associated with several diseases, most notably inflammatory bowel disease (IBD) (11, 12). However, stool samples alone do not fully reflect the total gut microbiota. Bacteria vary along the length of the gut as well as within specific niches such as the mucus layer overlying the intestinal epithelial cells (13). We and others have shown that the mucus microbiome is discrete from that detected in stools (14, 15) and that changes in the mucus microbiota precede changes in the stool microbiota and onset of disease (15). Host environmental factors such as diet and housing can also impact the microbiota, yet few studies report on how mice are caged and whether they are littermates (16-18). We have therefore aimed to test the effects of housing in our study.

To address and clarify some of the issues of potential inaccuracy and bias in microbiota analysis, we have developed a community-focused phylogenetic approach, without OTU calling, incorporating taxa at all phylogenetic scales. We use data from an IBD mouse model (colitis-prone, *mdr1a*^*-/-*^). These mice have been shown by standard analytical techniques to have an altered gut mucus bacteria (15). Importantly, the data come from a carefully controlled experiment, with co-housed *mdr1a*^*-/-*^ and wildtype mice. 16S sequencing was ordered to avoid confounding sequencing batches with treatments, enabling robust comparisons, among genotype, niche and age (15). To avoid taxonomic bias, we use the sequences themselves to estimate the phylogenetic relationships amongst organisms and thereby abundances at different phylogenetic scales. Only after analysis does our approach draw on a wider understanding of microbial taxonomy to interpret the findings. Our analysis use machine learning models to interrogate the microbiota of both stools and colonic mucus.

We find that the microbiomes of each cage are clearly distinguishable, being separated by differences in common, small-scale taxa (notably groups within the genus *Bacteroides*). Despite this variability, microbiomes from the different niches (stool and mucus) are clearly distinct, as are microbiomes from young (6-week) versus old (18 week) mice, each distinguished by taxa at intermediate phylogenetic and abundance scales. Surprisingly, despite the fact that our analyses are very effective at distinguishing these groupings and identifying the taxa involved, we were unable to find any consistent distinction between the microbiomes (stool or mucus), of *mdr1a*^*-/-*^ and wildtype (WT) mice at the stages tested. Thus, in removing potential experimental and analytical inaccuracies and biases, we identify, not only clear distinctions between microbial communities, but clear community robustness in the face of experimental perturbation.

## Materials and Methods

### Animal Maintenance

*Mdr1a*^*-/-*^ mice (FVB.129P2-Abcb1atm1Bor N7) (19) were bred with control FVB mice purchased from Taconic Farms (Albany, NY), to produce the F2 generation. Male mice from each litter were co-housed. Thus, WT and *mdr1a*^*-/-*^ mice from the same litters were used for all subsequent experiments. Male mice at 6 and 18 weeks of age were used for experiments. Food (Beekay Rat and Mouse Diet No1 pellets; B&K Universal, UK) and water were available *ad libitum*. Ambient temperature was maintained at 21 (+/- 2°C) and the relative humidity was 55 (+/- 10%) with a 12h light/dark cycle. All animals were kept under specific, pathogen-free (SPF) conditions at the University of Manchester and experiments were performed according to the regulations issued by the Home Office under amended ASPA, 2012.

### Isolation of genomic DNA

Sample collection and processing was performed as described by Glymenaki et al. (15). In brief, samples were harvested from mice at two time points, 6 and 18 weeks of age. Stool samples were collected from mice in individual autoclaved cages into sterile tubes and snap frozen on dry ice. Mice were sacrificed via CO2 inhalation, the proximal colon was cut open and the colonic mucus scraped using cell scrapers and Inhibitex buffer (QIAGEN, Manchester, UK) and snap frozen until use. Genomic DNA was extracted using QIAamp Fast Stool Mini-Kits according to the manufacturer’s instructions (QIAGEN). Additionally, snips of the proximal colon were fixed in Carnoy’s solution (60% methanol, 30% chloroform, 10% glacial acetic acid) and embedded in paraffin for histological analysis.

### Histology

Carnoy’s fixed colon samples were incubated in two changes of dry methanol (Sigma-Aldrich, Dorset, UK) for 30 minutes each, followed by absolute ethanol (ThermoFisher Scientific, Paisley, UK) for two incubations at 30 minutes each. Finally, tissue cassettes were processed in a Micro-spin Tissue Processor STP120 (ThermoFisher Scientific) and immersed in paraffin. Colon snips were embedded in paraffin blocks using a Leica Biosystems embedding station (Leica Biosystems, Milton Keynes, UK), with the luminal surface of the colon exposed for tissue sectioning. 5µm tissue sections were cut using a Leica Biosystems microtome and adhered to uncoated microscope slides (ThermoFisher Scientific). Slides were dried for 48 hours at 50ºC before use.

### Fluoresence *in-situ* hybridisation (FISH)

FISH was performed as described previously (15). In brief, FISH staining was performed using the universal bacterial probe-EUB338 (5′-Cy3-GCTGCCTCCCGTAGGAGT-3′), followed by immunostaining with a rabbit polyclonal MUC2 antibody and goat anti-rabbit Alexa-Fluor 488 antibody (Life Technologies, Paisley, United Kingdom). Slides were imaged using a BX51 upright microscope and a Coolsnap EZ camera (OLYMPUS, Tokyo, Japan) and images were processed using Image J.

### 16S rRNA gene sequencing processing

16S amplicon sequencing targeting the V3 and V4 variable regions of the 16S rRNA (341F: 5’-TCGTCGGCAGCGTCAGATGTGTATAAGAGACAGCCTACGGGNGGCWGCAG-3’ and 805R: 5’- GTCTCGTGGGCTCGGAGATGTGTATAAGAGACAGGACTACHVGGGTATCTAATC C-3’) was performed on the Illumina MiSeq platform (Illumina, California, USA) according to manufacturer’s guidelines and generated paired-end reads of 300bp in each direction. Illumina reads were demultiplexed to remove adapter sequences and trim primers. Illumina paired-end reads were merged together using SeqPrep (20) and submitted to MG-RAST’s metagenomics pipeline (21). Reads were pre-processed to remove low-quality and uninformative reads using SolexQA (22). The quality-filtering process included removal of reads with low quality ends (i.e. ambiguous leading/trailing bases) and the removal of reads with a read length two standard deviations below the mean. Artificial duplicate reads were then removed based on MG-RAST’s pipeline.

The resulting FASTQ files for every sample were merged into a single file of 590822 sequences to simplify processing, manually adding 3 known Archaeal 16S rRNA sequences from *Acidilobus saccharovorans, Sulfolobus tokodaii* and *Methanobrevibacter smithii*. Sequences were aligned using a specialist 16S RNA aligner using the Infernal algorithm (23), via a web-based interface provided by the Ribosomal Database Project (24). This file was then manually curated in R (25). Unless otherwise stated, all analyses were performed using custom scripts in R. The number of aligned bases in each sequence was recorded and the distribution of continuously aligned bases was examined. Any sequence that had less than 437 continuously aligned bases was discarded. The remaining 496550 sequences were taken forward for analysis. All sequences were identified using BLAST+ and the top hit for each sequence was recorded (26). The ‘classification’ function in the ‘taxize’ R package (27) was then used to assign full taxonomic information to each identified taxon where possible.

### Phylogenetic Tree

A phylogenetic tree of all sequences was generated using FastTree 2.1 (28), using the general time reversible (GTR) + CAT model and default parameters. The tree was rooted using the archaeal sequences as an outgroup. Phylogenetic clades were obtained using the ‘Ancestor’ function in the ‘phangorn’ R package (29). A relative abundance matrix, with abundance based on how many times sequences belonging to a phylogenetic clade appeared in a sample, was calculated.

### Ordination

Bray-Curtis dissimilarity values were calculated among all samples (based on the relative abundance matrix) and used for non-metric multidimensional scaling (NMDS) via the ‘MASS’ (30) and ‘ecodist’ R packages (31).

### Machine learning

Random forest (RF) models were run using the ‘randomForest’ package (32) in R. Specifically, the clade relative abundance matrix was used as an input for the RF, using a forest of 100,000 trees and the mtry value was left at default settings (the square root of the number of clades). Separate forests were run to predict whether a sample was 6 or 18 weeks old, whether a sample was stool or mucus, whether it was a WT or an *mdr1a*^*-/-*^ sample and what cage the sample was taken from. Each forest was controlled for all other treatments (i.e. a random forest comparing age included genotype and microbial niche as explanatory variables, in addition to the generated clades). The ‘MeanDecreaseAccuracy’ (MDA) value was used as a measure of how important each clade (or treatment) was at predicting treatment information and the out-of-bag (OOB) error rate was used to determine the predictive accuracy of the model. Nodes were ranked based on MDA value, taking the five most important nodes, determining the descendant tips and confirming the identity of the tip sequences via the BLAST+ results (26). Additionally, the depth of each node was determined using the ‘distances’ function in the igraph R package (33). A phylogenetic tree annotated with the resulting information was plotted using the ‘plot.phylo’ function in the ‘ape’ package (34).

### Validation of Model

In order to validate each model, we included a ‘randomised’ negative control RF where relative abundances of each node were permuted with respect to each sample and the predictive accuracy was assessed. In addition, we took the relative abundances of an important node for age and redistributed the abundance to only WT samples. The RF was repeated to investigate whether this node would appear as important for genotype. We also ran RF’s with an increasing number of trees, using three different random seeds and performed Spearman’s Rank correlation on the MDA values obtained among each set of three RFs of the same size. The Monod/Michaelis-Menten model was fitted, to determine how an increasing number of trees affected correlation of the MDA values. Finally, we included technical replicates of one stool sample that was used as an internal control between sequencing runs, in our forest models. We examined the MDA values for all the clades in each of these replicates to see how tightly correlated they were. Additionally, we compared the predictive accuracy of the RF model when using these different replicates.

### Statistical Analysis

Analysis of the real vs null RF models predictive accuracy was performed using 2-way ANOVA, with a Sidak’s post hoc test in GraphPad Prism 7 (GraphPad Software, La Jolla, USA).

## Results

### Phylogenetic tree of 16S rRNA data derived from the gut microbiota

Microbiota samples from the stools and colonic mucus of 40 male mice were collected at two different time points (6 vs 18 weeks of age) from two genotypes (WT vs *mdr1a*^*-/-*^) in an experiment using co-housed siblings of the different genotypes. Littermates, irrespective of genotype were co-housed in 7 cages. The colonic mucus resident bacteria were visualised by FISH (Figure 1a). Pathology was assessed and revealed that five of the 18 week-old *mdr1a*^*-/-*^ mice had indications of moderate or mild colitis, with a loss of healthy gut architecture and the remaining mice were healthy (15). Comparisons between healthy and colitic mice were tested in our subsequent experiments, but no consistent differences were identified (data not shown). On average, 10,442 16S sequences (range 1,892-25,681) were obtained per sample. All sequences were used to create a phylogenetic tree, comprising 496,550 tips and 428,234 internal nodes, which separated the major phyla Supplementary Figure S1). Sequences derived from stool and mucus were distributed across the tree (Figure 1b). Sequences associated with other criteria (age, genotype and cage) were similarly spread (Supplementary Figure S2).

**Figure 1:**
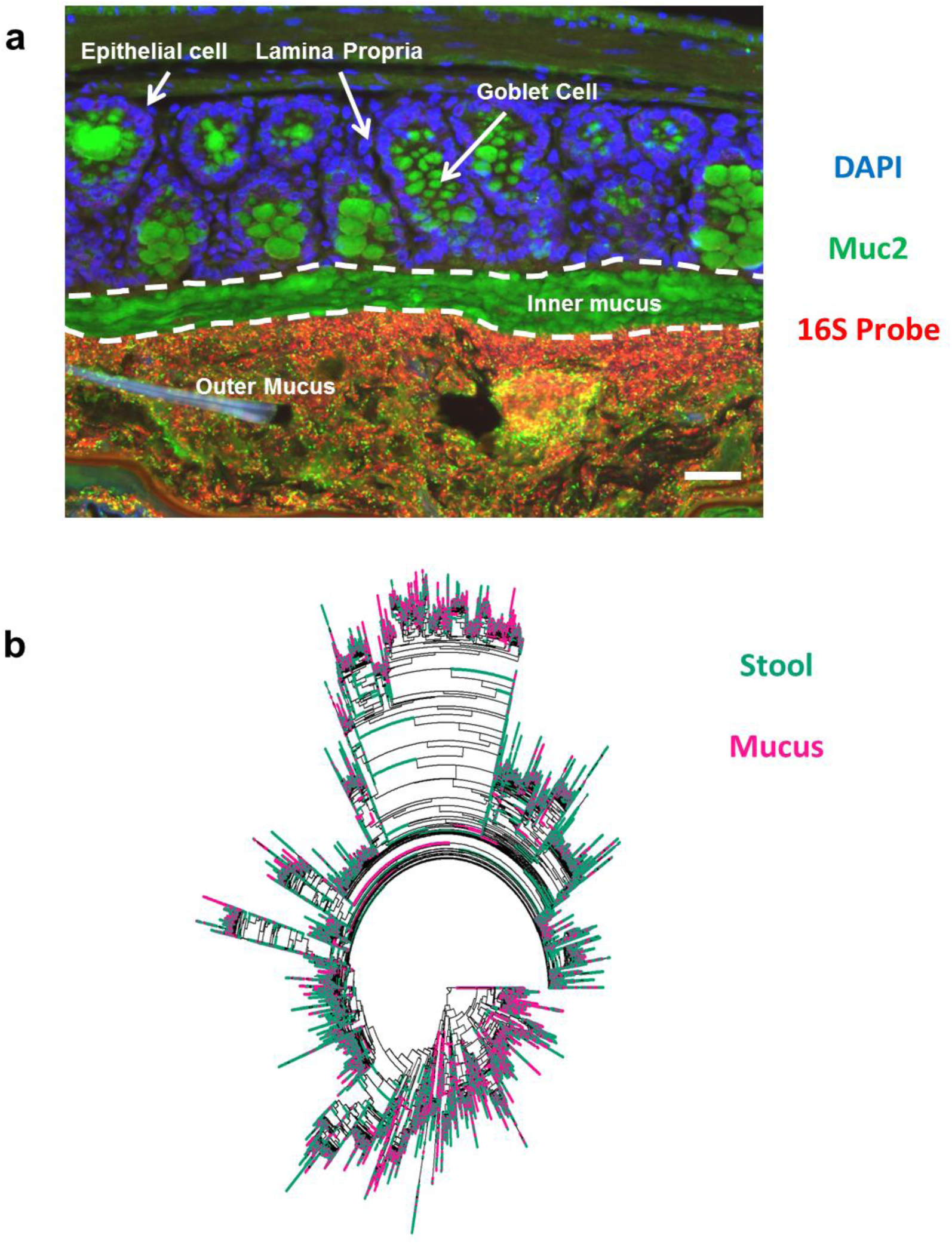
Distribution of stool and mucus-associated sequences across the phylogenetic tree. Colonic tissue sections from a male wildtype (WT) mouse was stained with a fluorescent DNA probe specific for the 16S rRNA gene to identify bacteria (red), a Muc2 antibody (green) to identify mucus and counterstained with DAPI (blue) (a). A phylogenetic tree of 16S rRNA sequences derived from the gut microbiota of FVB background WT mice and *mdr1a*^*-/-*^ mice (b). Colours indicate stool- (teal) and mucus- (pink) derived sequences, taken from n = 10-11 mice per genotype. The same tree is shown, coloured by other criteria, in Supplementary Figure S1 and S2.

### Strong separation of the gut microbiota by microbial niche, age and cage, but not host genotype

To avoid bias by taxonomic level, we constructed a data matrix comprising the relative abundance ((number of tips in clade in sample) / (total number of tips in sample)) of all 428,234 internal nodes of the phylogenetic tree in each of our samples. This avoided assigning OTUs, or using a reference database. To visualise the major differences in the microbial communities in an unsupervised fashion, Bray-Curtis dissimilarity values were calculated from our data matrix between all samples and used as an input for a 2-dimensional non-metric multidimensional scaling (NMDS) ordination (Figure 2). There was clear separation of samples by niche (Figure 2a). There was less separation by mouse age (6 vs 18 weeks) (Figure 2b) which is in concordance with our previous work (15). Samples from the same cage localised closely in the ordination. Cages containing different litters from the same mother were adjacent or overlapping, suggesting maternal effects influencing, but not fully explaining, cage-specific microbiomes (Figure 2c). Little separation was found when comparing genotypes (Figure 2d).

**Figure 2:**
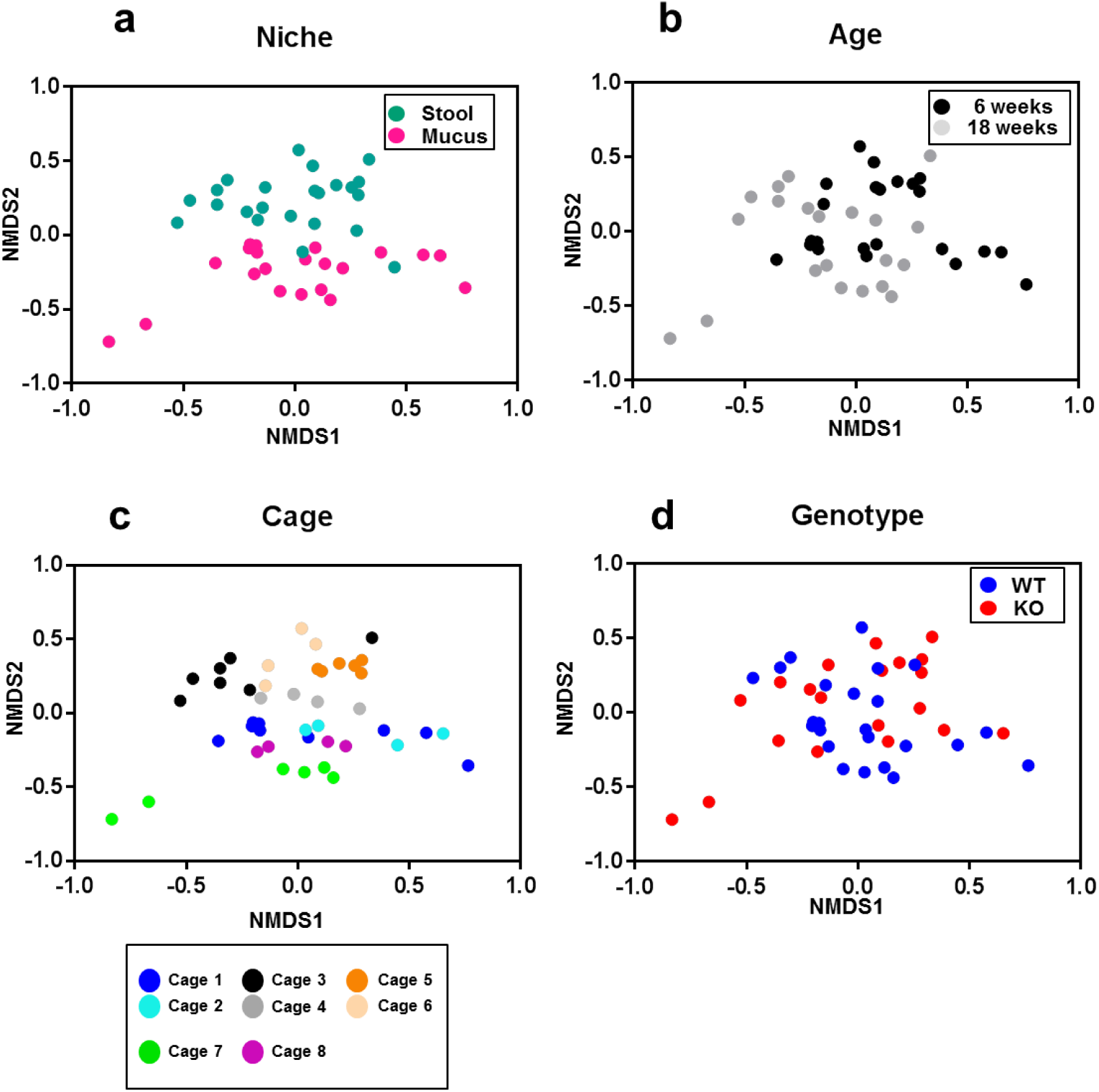
Separation of microbiota via NMDS for age, cage and microbial niche. Two dimensional non-metric multidimensional scaling (NMDS) was performed using a Bray Curtis dissimilarity matrix based on the relative abundance of all clades in the phylogenetic tree shown in Figure 1b. Plots highlighting 6 and 18 week samples (a), stool and mucus samples (b), different cages and mothers (c) and WT (wildtype) and KO (*mdr1a*^*-/-*^) samples (d) are illustrated. There were 5 mothers in total (cages 1 and 2, cages 3 and 4, cages 5 and 6, cage 7 and cage 8). Each point corresponds to a stool or a mucus sample. These samples were taken from n = 10-11 mice per genotype.

### Specific microbiota are strongly associated with age, microbial niche and cage but not host genotype

To determine the taxa driving the observed differences in community structure, the relative abundance matrix was used to construct machine learning models (random forests, RFs). Separate RF models were created to identify age, genotype, niche and cage based on the relative abundance of the clades (as defined by the phylogenetic tree, **Error! Reference source not found.**b) in each sample. These models were compared against a null (negative control) model where relative abundances were permuted among taxa within samples to remove true associations. Niche could be determined from the microbiota with 92% accuracy, age with 98% accuracy and cage with 80% accuracy (averaged across six technical replicates), in all instances substantially higher than the negative control model (**Error! Reference source not found.**a) (Two Way ANOVA- Sidak’s post hoc test: *P* < 0.0001). Genotype could not be determined from the microbiota using our RF models any better than in the negative control (Figure 3a). Models considering genotype will therefore not be considered further.

**Figure 3:**
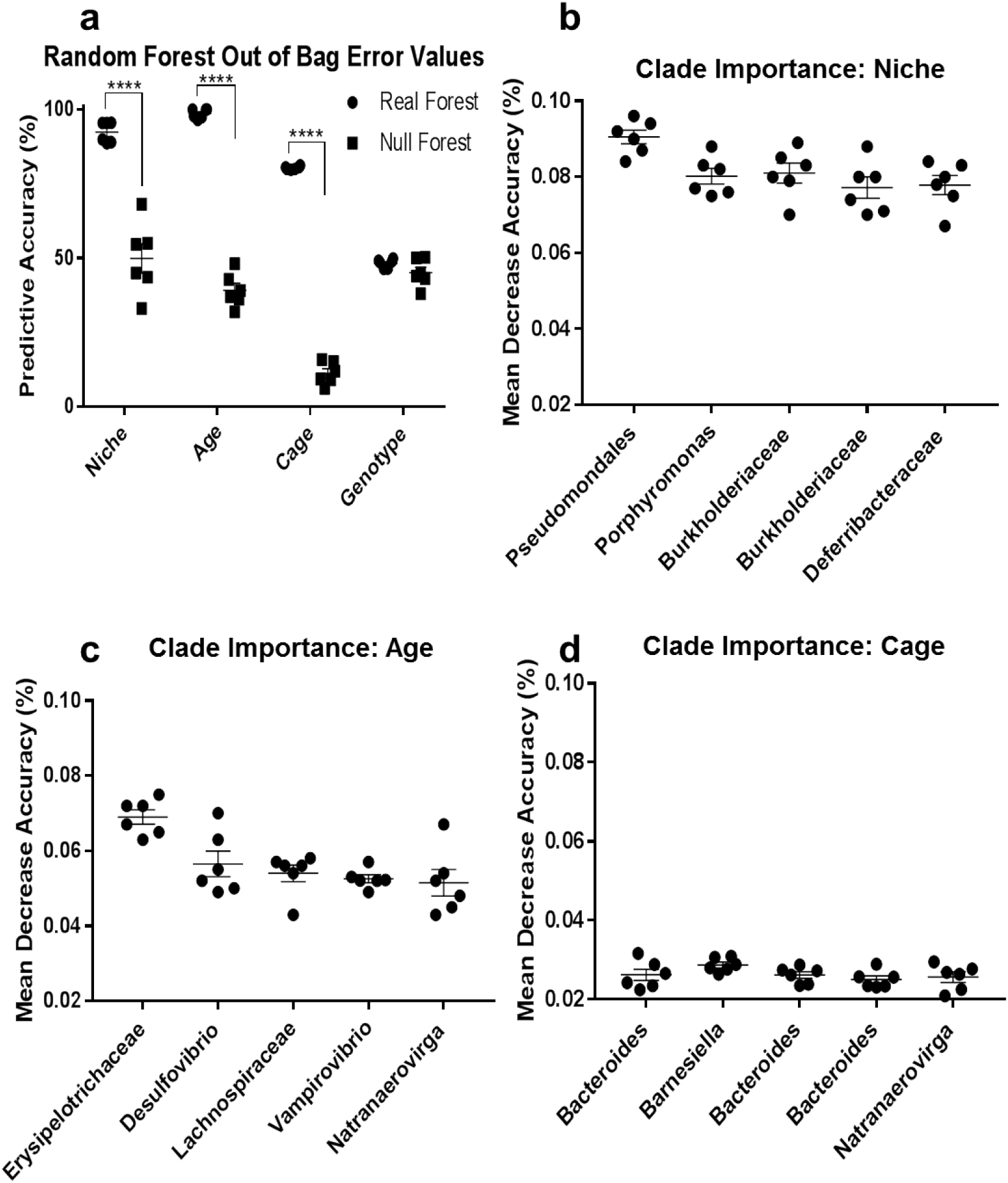
Random forest model identifies strong associations between the microbiota, niche, age and cage. The predictive accuracy of the random forest model at taking a sample and discriminating between the different treatment groups is shown (a). The five most important nodes associated with microbial niche (b), host age (c) and social group (cage) (d), are named based on the finest scale taxon containing the closest BLAST hits of all sequences in the clade. Bars represent means and standard errors. Asterisks represent significance determined using Two Way ANOVA-Sidak’s post hoc: *P* < 0.00001 (****). Similar plots for further validation control analyses are shown in Supplementary Figure S3 and S4. n = 6 technical replicates.

The RFs give an importance value for each clade (the tree’s internal nodes) in discriminating between groups. To identify which bacteria the clades encompassed, we used BLAST+ on all sequences, recording the taxonomic identity of the top hit (hits that had a percentage coverage <100% were discarded). The finest-scale taxonomic grouping containing all sequences descending from the five most important clades is shown in Figure 3b, c and d for niche, age and cage RFs respectively.

For microbial niche, the most important distinguishing clades were all Gram negative and mostly comprised *Proteobacteria*: the order *Pseudomonadales* and three clades in the families *Burkholderiaceae* and *Deferribacteraceae* and all these clades were more abundant in the mucus samples. The *Deferribacteraceae* containing clade mostly comprised *Mucispirillum*, a known mucus-associated bacteria (35). The genus *Porphyromonas* was more abundant in stool samples.

The most important clades separating ages were the families *Erysipelotrichaceae* and *Lachnospiraceae* within the *Firmicutes* phylum (which have each been specifically associated with young mouse microbiomes before (36)) plus three genera: *Natranaerovirga Desulfovibrio*, and *Vampirovibrio* in the *Firmicutes, Proteobacteria* and *Cyanobacteria* phyla respectively. With the exception of *Natranaerovirga*, all these bacteria were prevalent in the 18 week old mice.

The most important clades separating cages were *Natranaerovirga* (a different clade from that important for separating ages) plus four clades within the order *Bacteroidales* – three comprising the genera *Bacteroides* and one comprising *Barnesiella*.

To validate the method for identifying important taxa, we firstly tested the reproducibility of different estimates of clade importance based on the same data. We expect more reproducible estimates (i.e. tighter correlations among estimates) when more trees are used in the random forest, which we find to be true (Supplementary Figure S3a). We also expect there to be a maximum possible correlation among independent importance estimates. By calculating the rank correlation among importance values from independent RFs of different sizes and fitting a saturating function (Supplementary Figure S3b), we estimate that a forest of 100,000 trees, as we use in our analyses, achieves 99% of the maximum correlation of 0.21; i.e. the approach used in the present study gets close to optimal reproducibility. Secondly, we validated the approach to individual importance estimates by taking the most important clade identified for age (within the *Erysipelotrichaceae* family) and redistributing the abundances of this clade to only be present in all WT samples. Upon repeating the random forest separating on genotype, the *Erysipelotrichaceae* clade became the most important and this change meant the genotype RF was now significantly better than the null model (Two Way ANOVA-Sidak’s post hoc test: *P* < 0.05) (Supplementary Figure S4a). Similarly, repeating the random forest separating on age using the redistributed dataset, the *Erysipelotrichaceae* clade ceased to be the most important, leaving most of the other important clades largely unaffected (Supplementary Figure S4b). This suggests that important clades are robust and maintain their importance, even when other clades are altered. Thirdly, we included technical replicates of one stool sample that was used as an internal control between sequencing runs, in our forest models. The results of these technical replicates were highly correlated (Supplementary Figure S5a) and which technical replicate was included made little difference to the results (Supplementary Figure S5b).

### Abundant, low-level taxa distinguish cage microbiomes but not age or niche

Having identified taxa at different phylogenetic levels as particularly important for separating microbiomes, we looked systematically at the phylogenetic scales that are important for separating different microbiomes. Clade importance was analysed as a function of the number of nodes between the clade and the root of the phylogenetic tree (Figure 4) or the distance from each clade to the root (Supplementary Figure S6). These measures distinguish between clades with fewer nodes and shorter branch lengths between them and the root (high-level taxa) corresponding, e.g. to phyla, and clades with more nodes and longer branch lengths between them and the root (low-level taxa) corresponding e.g. to genera. For both age and niche, neither the lowest nor the highest level clades were important (little separation from the null model), but the important clades were of intermediate taxonomic levels (Figure 4 a-b). However, for differences among cages, while intermediate level clades were important, many of the most important groups were at the extreme of low level taxa, i.e. differences in sub-specific groupings (Figure 4c, Supplementary Figure S6c).

**Figure 4:**
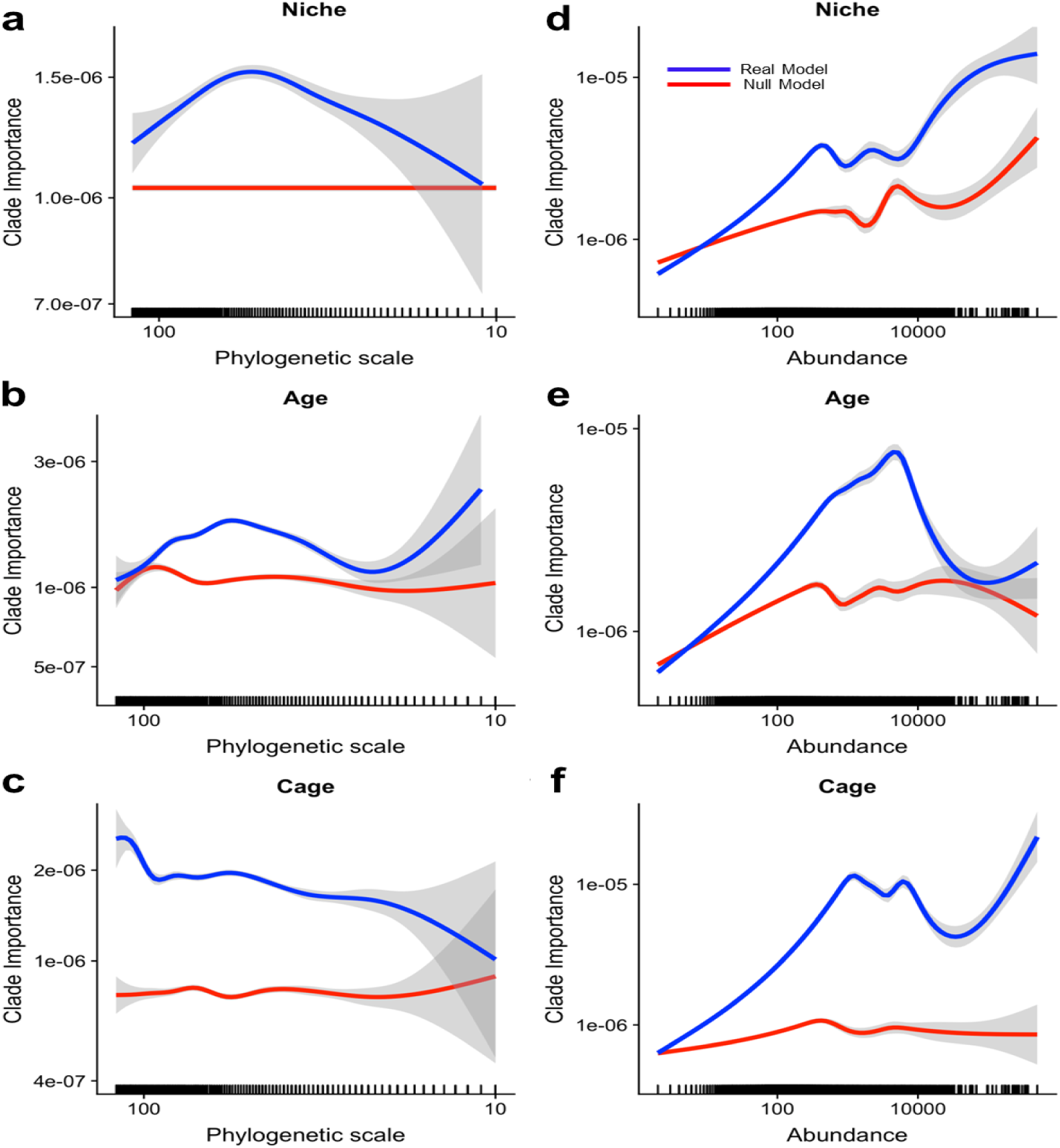
Abundant, low-level taxa distinguish cage microbiomes but not age or niche. The phylogenetic scale of each clade was measured as the number of nodes in the phylogenetic tree between the clade and root (small values associated with large-scale taxa such as phyla). Phylogenetic scale was compared against the ‘mean decrease in accuracy’ (MDA) value when running a forest that distinguished the niche (a), age (b) and cage (c). Clade abundance was measured as the number of sequences (tips) descending from each clade. Each clade’s abundance was compared against its MDA value, when running a forest that distinguished the niche (d), age (e) and cage (f). The smoothed mean for the ‘real’ random forest model is illustrated in blue and for a null (negative control) random forest model in red. The grey areas refer to confidence intervals. Note the logarithmic scales on all axes. n = 6 technical replicates. Similar plots using a different measure of phylogenetic scale (distance of clades from the root of the tree) are in Supplementary Figure S6.

The number of sequences within a clade of the tree that were present in a particular set of samples is an estimate of its abundance in that microbiome. We therefore asked how abundance of bacterial taxa correlated with its importance in distinguishing microbiomes. We found that, for separating age, niche or cage microbiomes, moderately abundant taxa were important, whereas the rarest taxa were never important (Figure 4d-f). The most abundant taxa were important for distinguishing cage microbiomes, but were much closer to the null expectation for distinguishing microbiomes from different ages or niches.

The importance of particular taxa was uncorrelated across forests that distinguished different criteria (age versus niche, age versus cage and niche versus cage; rank correlation −0.002, 0.005 and −0.001 respectively). There was also a large overlap between distributions of clade importances between true and null forests (Supplementary Figure S7). Thus, while we can identify broad trends (Figure 4) and a few key taxa associated with particular gut features (Figure 3), we did not find widely applicable ‘indicator’ taxa that were individually sensitive to multiple effects on the gut microbiome.

## Discussion

Our modelling approach discriminated clearly between the microbiomes of 6 and 18 week old mice, between mucus and stool samples and between different groups of co-housed mice. This confirms the robustness of our model as it is consistent with others’ work, for instance showing microbiome changes with age in both humans and mice (37, 38) and work identifying microbial niche as the strongest factor for separation of the microbiota (15, 39). Our mice are still relatively young, initial samples taken only ∼2 weeks after weaning and at 18 weeks old. Therefore the differences seen by age may be due to the microbiota adjusting due to changed diet. Nonetheless, microbial changes in mice can happen within as little as two weeks (40). The fact that we saw these differences clearly and identified the key taxa responsible, validates our approach to modelling these microbiome changes. It is therefore striking that even this apparently powerful approach could find no consistent differences between the gut microbiomes of co-housed wildtype and colitis-prone (*mdr1a*^*-/-*^) genotypes.

Differences in the microbiota of WT and *mdr1a*^*-/-*^ mice *have* been reported (15, 41). The fact that we do not see them here (Figures 2d and 3a) is therefore unexpected. Discrepancies in sample size between treatment groups can be a problem for RFs applied to such data (42). However, here sample sizes are well balanced (20 wildtype and 20 *mdr1a*^*-/-*^ mice). Some, but not all, of the older *mdr1a*^*-/-*^ mice were starting to develop colitis. Therefore, changes in the microbiome with the onset of colitis may have obscured any consistent differences among genotypes. It may also be that any machine learning approach used on such a dataset would be under-powered with too small a sample size to identify subtle differences. Alternatively, previous analyses may have been misled by large cage effects (Figures 2c and 3a) into erroneously attributing some of that variation to differences among genotypes. Regardless, it is clear that, in comparison to differences between gut age and niche, the mouse gut microbiome is relatively robust to this host genotype change affecting the gut. This is consistent with previous findings where the effect of host environment on microbial communities has been much greater than that of host genetic effects (16).

The majority of the gut microbiota fall within a small range of phyla, with *Firmicutes* and *Bacteroidetes* making up the largest proportion and *Actinobacteria, Proteobacteria, Fusobacteria* and *Verrucomicrobia* a smaller proportion (Supplementary Figure S1, (43)). Shifts in the proportions of phyla have been associated with physiological effects on the host, for example increased Firmicutes is associated with increased incidence of obesity (44, 45). In the context of IBD, numerous phylum level changes have been associated with the progression of inflammation. A reduction in the abundance and diversity of *Firmicutes* is associated with IBD in human patients (46-48) and *Bacteroidetes* has been shown to be both increased (49) and decreased (46). However, our data did not find such high-level taxa to show consistent differences in any of our microbiome comparisons (Figure 4a-c, Supplementary Figure S6). These high-level taxa are also abundant taxa and, *a priori*, it might have been reasonable to expect that the more abundant taxa would have the most important functional consequences for the host and therefore be most likely to differ between different circumstances. However, the only microbiome comparison in which we find the most abundant taxa to be important was in distinguishing among cages and even here, it is low-level taxonomic groupings (e.g. clades within the abundant genus *Bacteroides*), not phyla that distinguished among cage-specific microbiomes.

Rare species have been suggested to play a large role a range of ecosystems, including the host and the environment (50, 51) and rare taxa have been associated with inflammation (52). We previously showed that lower level taxonomic changes can also have functional significance, with the genus *Pseudomonas* causing a delay in wound repair (53). Our analysis here however did not find the rarest taxa to be important in discriminating between microbiomes. This could be an artefact of the fact that, almost by definition in a complex microbiome, rare taxa are likely to be missed from at least a subset of samples through random sampling. Therefore, rare taxa would not show consistent differences among the factors considered (age, niche or cage), as we find (Figure 4d-f).

Several specific taxa we highlighted as being important in distinguishing microbiomes (Figure 3b-d) included taxa that have been associated with differences in gut microbiomes before. This helps validate our analytical approach. We implicated the family *Erysipelotrichaceae* in distinguishing older mice, a subset of which are colitis-prone compared to younger healthy mice. *Erysipelotrichaceae* have been associated with the development of IBD in an infection-induced mouse model of colitis (54) as well as colorectal cancer which is known to be a potential risk of IBD (55). However, this bacterial family has also been reported as significantly decreased in a murine model of colitis driven by tumour necrosis factor (TNF) (56). It will be important, given the impact of cage and maternal effect, that we carefully re-consider previously published data. Notably, the presence or absence of certain taxa will allow other bacterial families/species to flourish or be inhibited, which in turn will alter host/microbial homeostasis, emphasising the need to consider communities and not bacteria in isolation. Furthermore, changes in one bacterium may not be significant functionally if the clade as a whole is unaffected. Our RF models can account for interactions among taxa, and the ‘importance’ assigned to a taxon takes these into account (57).

In distinguishing between microbial stool versus mucus niches (Figure 3b), we did not identify typical lumen/faecal associated bacteria such as *Ruminococcus*, as important for distinguishing niche (58). The most important clade identified was one that encompassed bacteria within the order *Pseudomonadales*. This order includes genera such as *Pseudomonas and Acinetobacter. Acinetobacter* species are known to be associated with the colonic mucus (59) and therefore likely to be good markers of the mucus microbiota. The family *Deferribactericeae* also distinguished mucus and stool and this family contains the genus *Mucispirillum*, known mucus-associated bacteria (60). Again, these findings validate our approach, showing it capable of identifying particular taxa that distinguish gut niches in a more nuanced way than traditional correlative methods. However, it was notable that a common mucus-degrading bacteria, *Akkermansia muciniphila*, (61) was not found to be important. This may be because its occurrence is more variable among treatment groups and indeed our previous work suggested *Akkermansia muciniphila* only became prevalent in the mucus of inflamed guts (15). Again this emphasises the need for caution when interpreting apparent microbe-niche associations using single species level analysis.

RF models can be used to address clear questions about the microbiome, while also taking account of its complexity. For example, RF could discriminate between lean and obese subjects, where simple summary statistics such as the ratio of Firmicutes to Bacteroidetes could not (42). Similarly, RF was used to discriminate between patients with active Crohn’s disease and those in remission with ∼70% accuracy (62). By building a phylogenetic tree of the sequences and using the full range of clades in that tree as explanatory variables in the RF model, we can identify particular clades as important, whatever taxonomic level they occur at. All clades are treated in an equivalent, data-driven way and we can ask what big-scale patterns exist in the relative importance of clades at different scales and abundances (Figure 4, Supplementary Figure S6). A risk of our approach is losing the connection to specific microbial taxa. However, simple post-hoc similarity searches of the sequences involved were effective at naming key taxa involved (Figure 3b-d), demonstrating both the similarities, such as the importance of *Natranaerovirga* in distinguishing both cages and ages, and differences. For example, although similar phylogenetic levels and taxon abundances distinguished niches and ages (Figure 4a-b, d-e), Proteobacteria clades were most important for niche and Firmicutes clades for age (Figure 3b vs. 3c).

Building phylogenetic trees only using the information in amplicons from subsets of the 16S rRNA (Figure 1b) is restrictive. Even 16S-based trees using the complete sequence from carefully chosen bacteria do not fully capture their evolutionary history (63), and our tree therefore does not fully capture the topology of more thorough 16S-based trees (64). Agreement with accepted evolutionary trees is only possible for such partial 16S sequences by incorporating many constraints (as done, e.g. by Louca et al. (65)). Nonetheless, taking this restrictive approach ensures that the power of the data is fully used without accepting the biases that such constraints undoubtedly impose by attempting to shoe-horn data into a pre-existing framework. The approach described in this paper avoids the risks both of over-stretching the data such as assigning a sequence read to one taxon rather than another when it is in fact similarly very close or very distant to both. It also ensures that we do not lose power that is in the data e.g. clear phylogenetic structure among sequences that are closer than a given threshold, typically 97% identity used for OTUs.

The development of ‘de-noising’ approaches such as DADA2 (66) and DEBLUR (67) to generate amplicon sequence variants (ASVs) also goes some way to avoiding the problems of using universal similarity thresholds to define OTUs. Our approach goes one step further, and avoids issues of de-noiser choice (68), by using all sequence variants, whether true ASVs or sequencing errors. Given our well-controlled experiment, we do not expect different sequencing errors in different treatments. This is validated by the fact that we find the very rarest variants, which will be highly enriched for sequencing errors, are no better than random at distinguishing any of our treatments (Figure 2d-f). Our focus on differences among treatments comes at a cost – we do not even attempt to estimate the ‘true’ community composition of any particular sample. Despite this, we are able to identify clear compositional differences in communities across the treatments studied.

In conclusion, taking a carefully designed factorial experiment involving co-housing of different mice with genotypes that affect their susceptibility to IBD, we have been able to identify major changes in the gut microbiome with age, niches and cages, but not genotype (Figures 2, Figure 3a). Our machine learning approach, focused on phylogenetically related groups at all evolutionary scales, proved effective, not only in identifying distinct versus homeostatic microbiomes, but in identifying the phylogenetic groupings important in making distinctions (Figure 3b-d). Our approach has therefore gone beyond investigating single species changes. Furthermore, this approach revealed differences in the patterns of phylogenetic groupings (high or low level, rare or abundant taxa) that distinguish different microbiome features (Figure 4). Together, this work reveals the subtlety of the balance between homeostasis and difference in the gut microbiome, that can be used to better define the host interaction with its microbiome, and can be applied to many and diverse conditions to better understand the role of the microbiome in health and disease.

## Declarations

### Acknowledgements

We would like to thank Dr Maria Glymenaki for sample collection in the initial study. We would also like to acknowledge the Histology, Genomics and the Computational Shared Facility at the University of Manchester and funding from the BBSRC for a DTP studentship for GS and funding from The European Crohn’s and Colitis Organisation (ECCO) awarded to SC.

## Competing interests

The authors declare that they have no competing interests.

## Availability of data and material

The sequence data analysed during the current study is available on the European Bioinformatics Institute (EBI, https://www.ebi.ac.uk/ena) (study accession number PRJEB6905). Code to reproduce the main text figures produced in R will be uploaded to GitHub.

