## Supplementary Figures for "Distinct microbial communities in the murine gut are revealed by taxonomy-independent phylogenetic random forests"

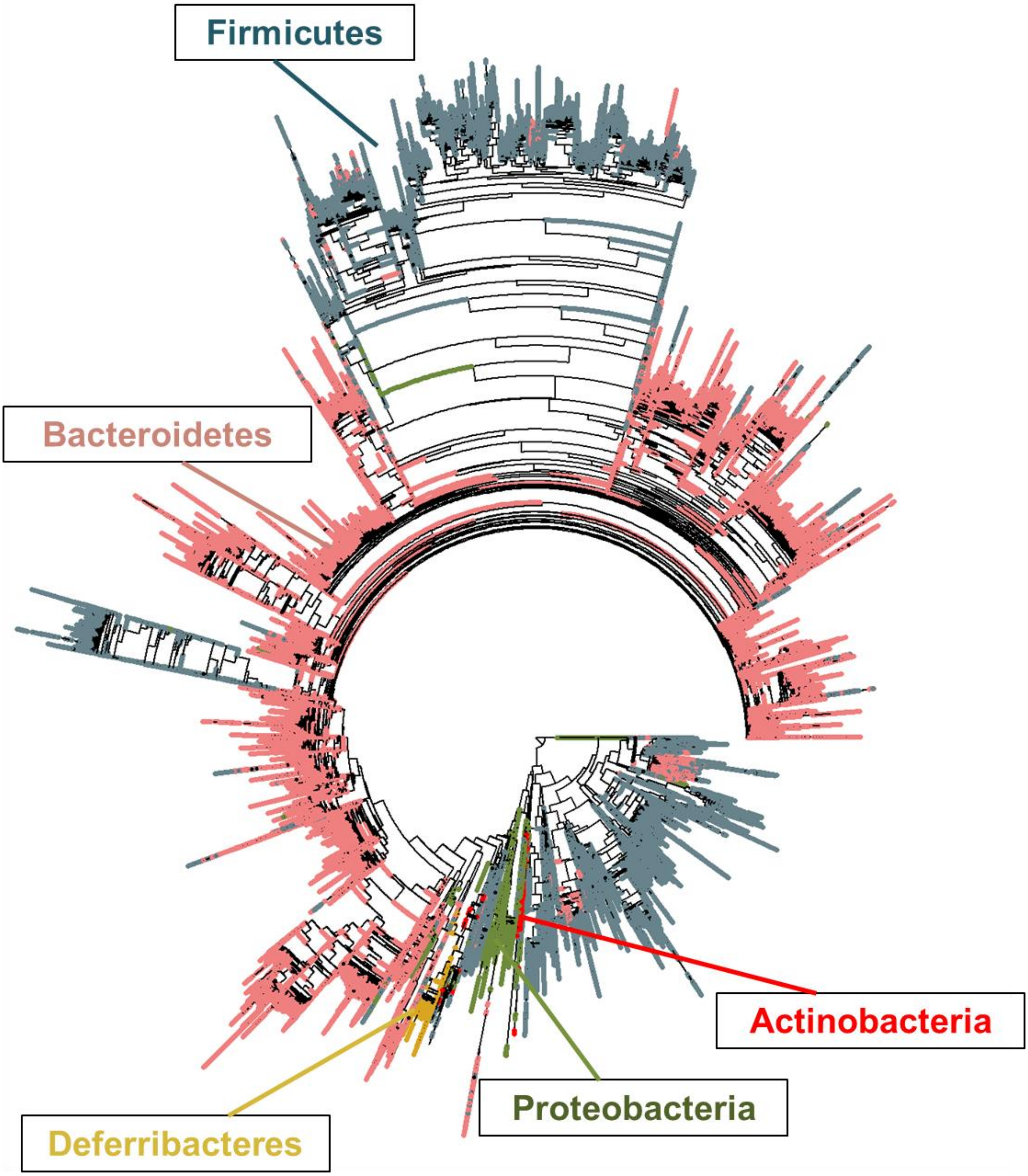


**Supplementary Figure S1: Major phyla are represented on the phylogenetic tree.** A phylogenetic tree of 16S rRNA sequences derived from the gut microbiota of FVB wildtype (WT) mice and *mdr1a*$^{-/-}$ mice. The distribution of major gut phyla are highlighted on the tree: Firmicutes (grey), Bacteroidetes (pink), Proteobacteria (olive), Actinobacteria (red) and Deferribacteres (gold).

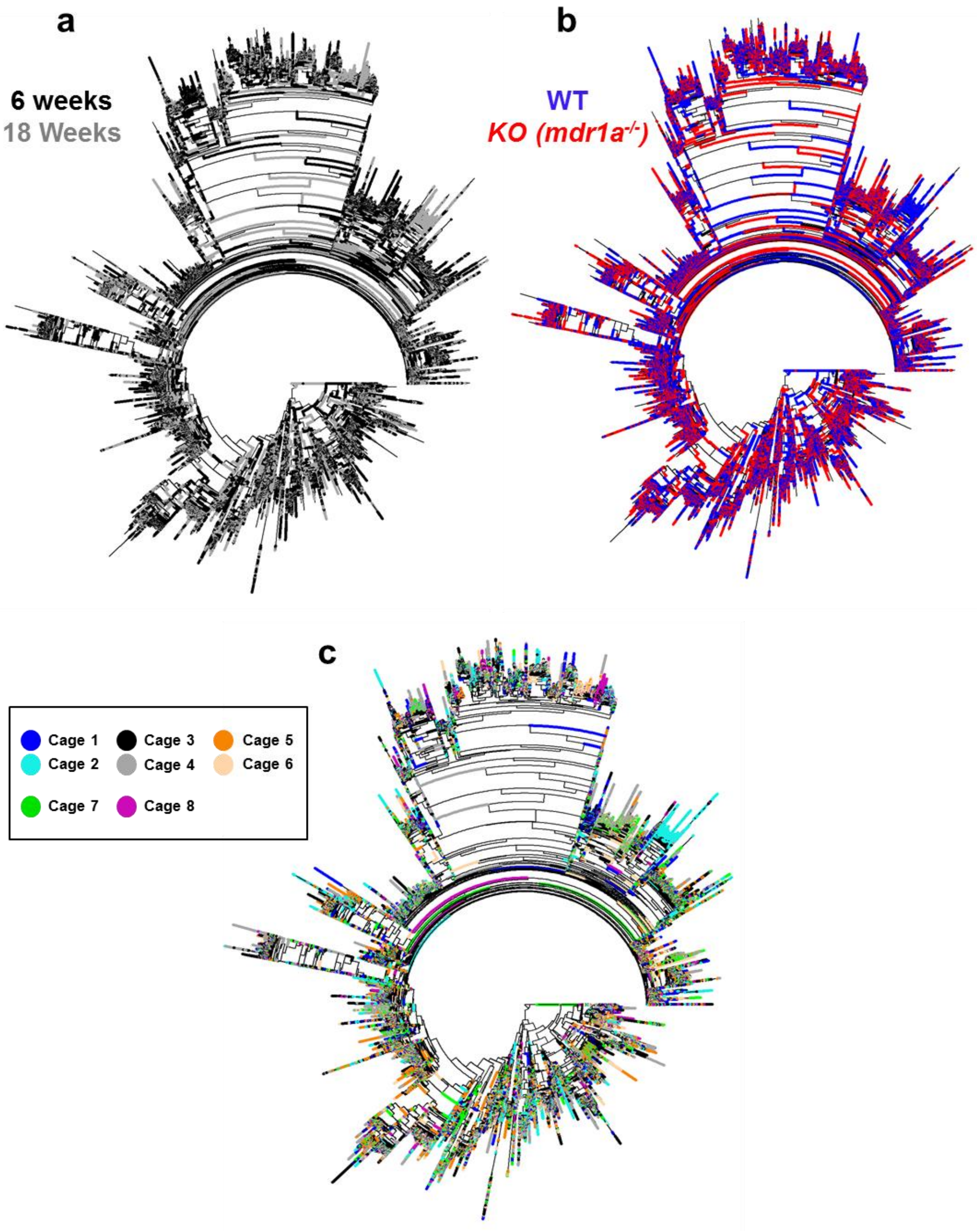


**Supplementary Figure S2: Wide distribution of sequences on a phylogenetic tree when coloured by treatment groups.** A phylogenetic tree of 16S rRNA sequences derived from the gut microbiota of FVB wildtype (WT) mice and *mdr1a*$^{-/-}$ mice was plotted and coloured by: Age (a), genotype (b) and cage (c). Colours indicate 6 weeks of age (black), 18 weeks of age (grey), WT mice (blue), *mdr1a*$^{-/-}$ mice (red) and different cages (red, green, goldenrod, purple, dark blue, steel blue, pink and dark green).

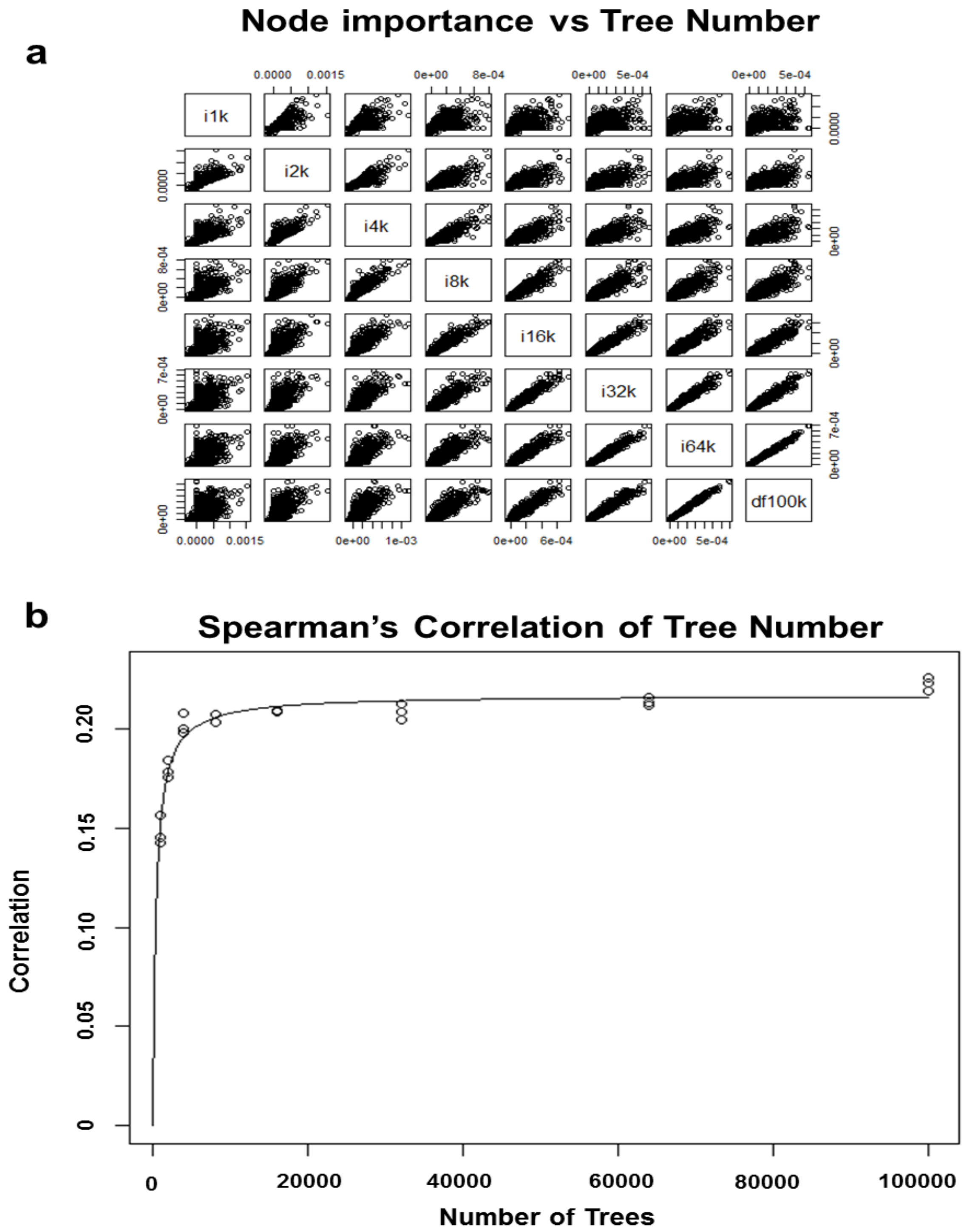


**Supplementary Figure S3: Assessing Robustness of the Random Forest (RF) model.** A RF for separating stool and mucus was run using a particular random seed with an increasing number of trees in R and the 'MeanDecreaseAccuracy (MDA) value was plotted for each clade in all forests (a). Spearman's rank correlation between clade importance in three RFs separating stool and mucus, run with different random seeds and a particular number of trees. The best-fitting saturating (a Monod/Michaelis-Menten) curve is also shown (*Max* = 0.21, *K* = 439.97) (b).

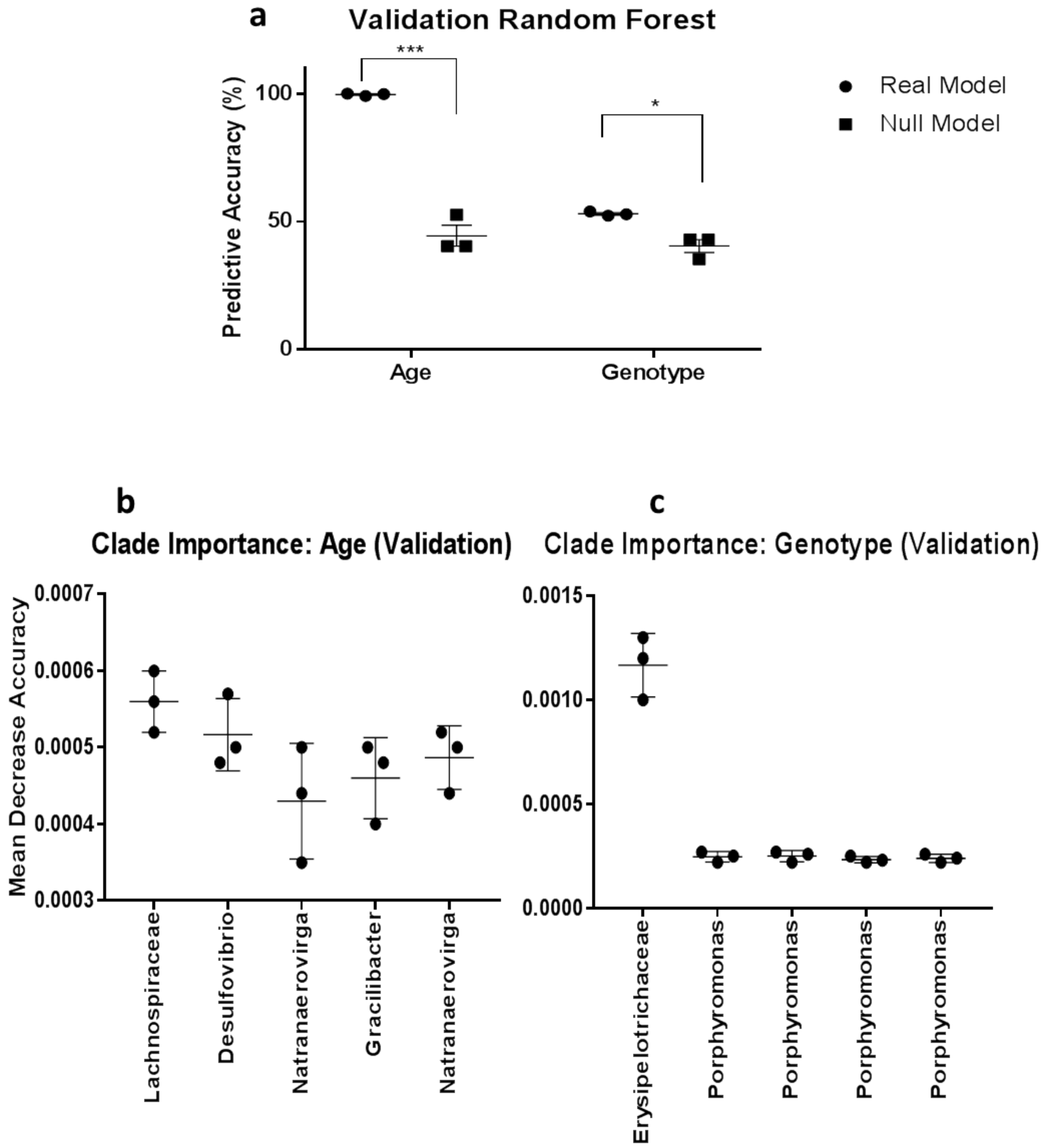


**Supplementary Figure S4: Redistributing the relative abundance of clade Erysipelotrichaceae confers importance to genotype.** The most important clade for age was identified and its relative abundances were redistributed into wildtype samples only and a random forest (RF) was run. The predictive accuracy of the RF model at taking a sample and discriminating between age and genotype are displayed (a). The five most important nodes when comparing age (b) and genotype via the RF are illustrated. Data shown as mean +/- standard error mean (SEM). Asterisks represent significance determined using Two Way ANOVA: $p =< 0.00001$ (****). n = 3 technical replicates.

a

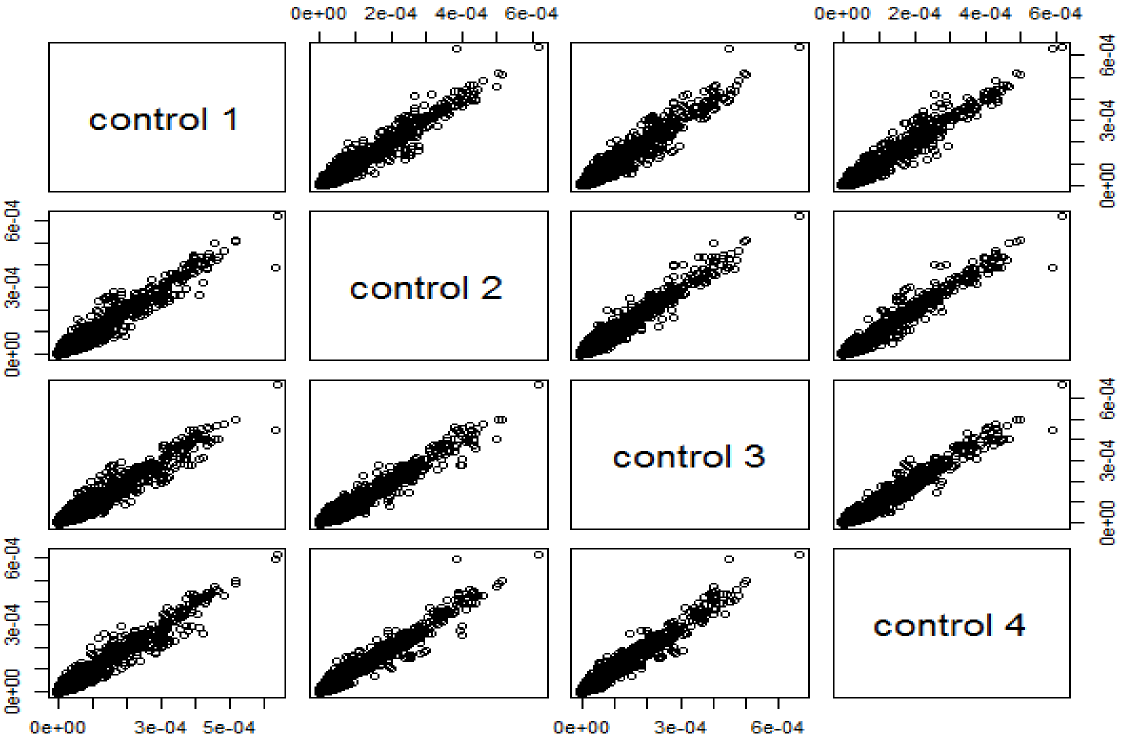


b

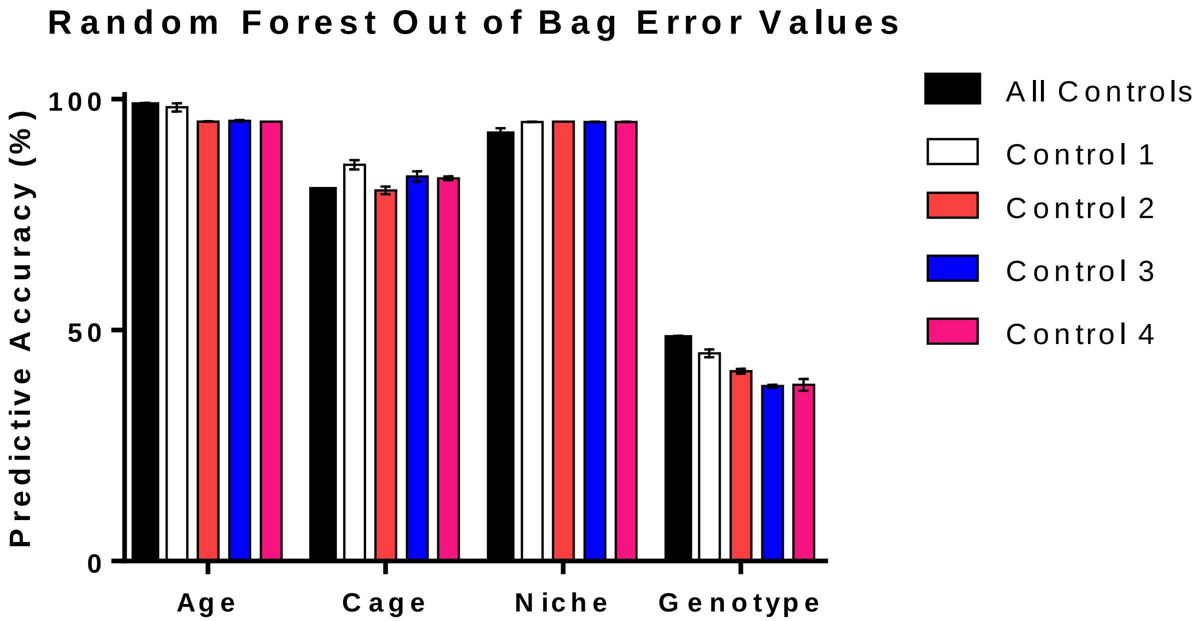


**Supplementary Figure S5: Control samples are highly correlated.** 16S rRNA was sequenced from the stools and mucus, of 6 and 18 week old, male wildtype (FVB background) and *mdr1a*$^{-/-}$ mice. One 'control' stool sample was sequenced multiple times to account for variability between sequencing runs. A random forest (RF) was performed excluding all but one of these control samples and the correlation of 'mean decrease accuracy' values was plotted (a). The predictive accuracy when running a RF forest including all these controls or excluding all but one was plotted (b), for each of the treatment groups (age, cage, niche and genotype).

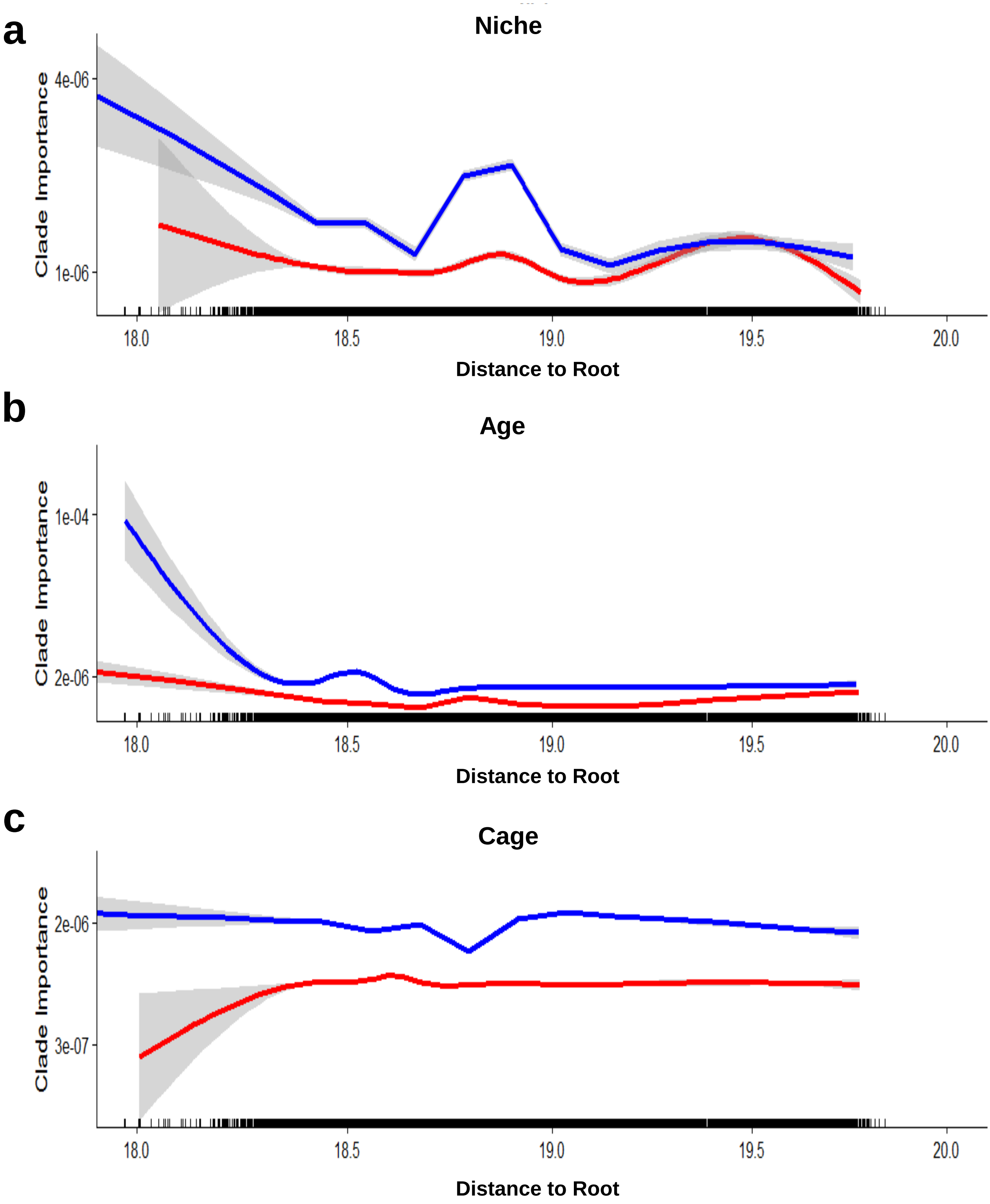


**Supplementary Figure S6: Taxa of intermediate level distinguish niche, age and cage.** The distance from clade to root was compared against the 'mean decrease accuracy' (MDA) value when running a random forest (RF) that compared the niche (a), age (b) and cage (c). The 'real' RF model is illustrated in blue and a null (negative control) RF model is illustrated in red.

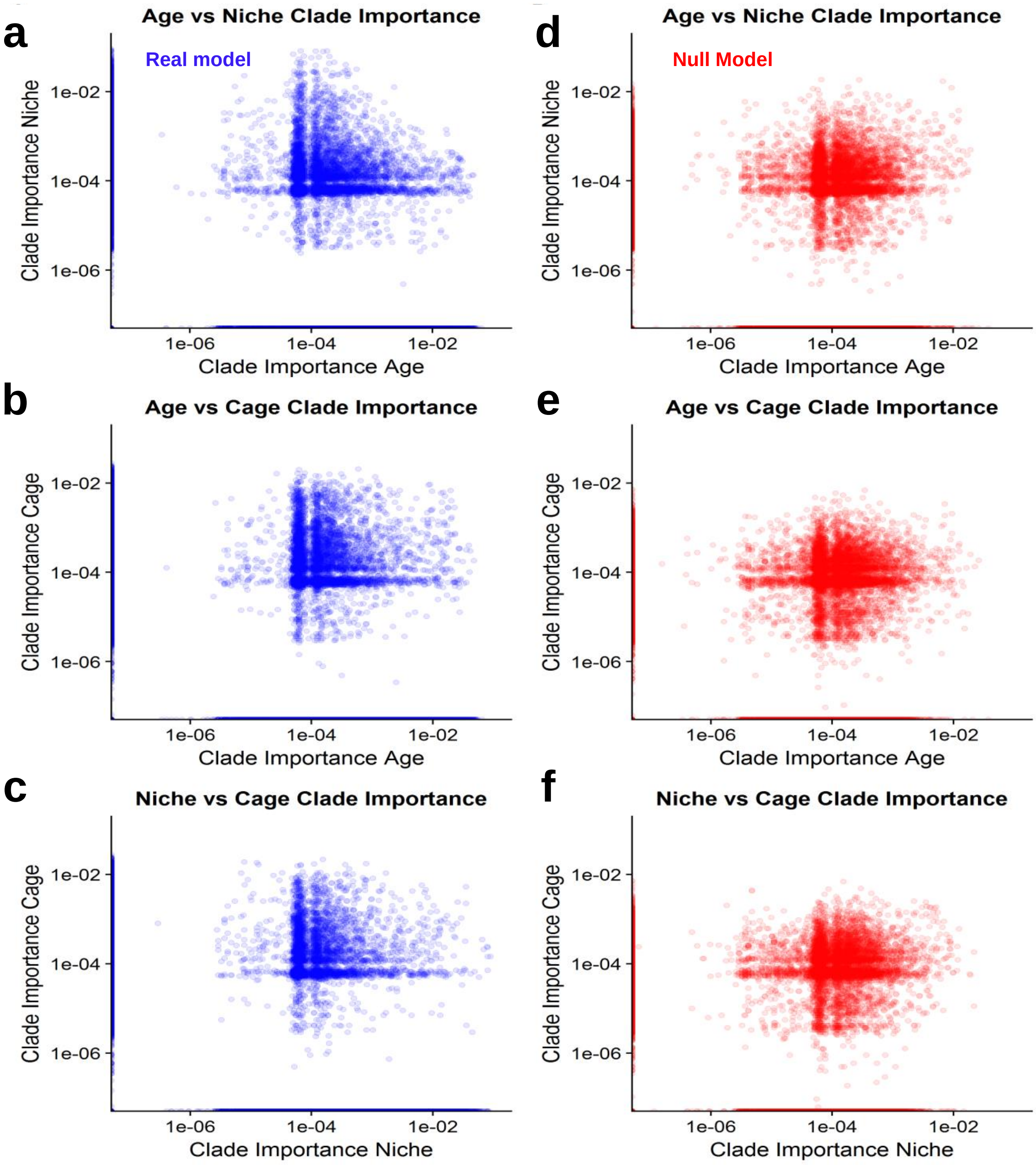


**Supplementary Figure S7: Clade importance between forests.** The importance (Mean Decrease Accuracy value) of a clade in one random forest (RF) was plotted against its importance in another RF for the real model (blue): Age vs Niche (a), Age vs Cage (b), Niche vs Cage (c) and the null (negative control, red) model: Age vs Niche (d), Age vs Cage (e) and Niche vs Cage (f).
